# TFAP2A links drug resistance to antitumor immunity

**DOI:** 10.64898/2026.07.08.735861

**Authors:** Haiwei Mou, Veronika Yakovishina, Kristen DeRosa, Yeqing Chen, Min Xiao, Maggie Dunne, Nancy Shi, Monzy Thomas, Jordan L. Smith, Qin Liu, Meenhard Herlyn

## Abstract

Combination targeted therapy with BRAF/MEK inhibitors and immune therapy show promising therapeutic outcomes in melanoma; however, the development of drug resistance still represents a formidable challenge. Remaining unexplored is the possibility that BRAF/MEK inhibitors themselves inadvertently compromise the tumor immune microenvironment, limiting the efficacy of immunotherapy when it is used in combination with targeted inhibitors. Herein, we profiled the landscape of the BRAF regulatome identifying a novel transcription factor, TFAP2A, newly linking BRAF/MEK drug resistance to antitumor immunity. Specifically, we found that BRAF/MEK inhibitors significantly upregulate TFAP2A. Further, genetic disruption of TFAP2A overcomes BRAF/MEK-inhibitor resistance, promotes stromal enrichment, and enhances intratumoral infiltration of macrophages in an immune-compromised mouse model. In a syngeneic mouse model, TFAP2a knockout not only suppresses tumor growth but also induces potent anti-tumor tertiary lymphoid structures (TLSs). Single cell transcriptomics revealed that the absence of TFAP2A shapes the antitumor microenvironment with an influx of M1-like macrophages, CD8+ T cells and mature dendritic cells. By identifying TFAP2A as a shared driver of both targeted therapy resistance and immunosuppression, our work offers a one-stone-two-bird strategy to overcome drug resistance and elicit antitumor immunity.

## Introduction

Although targeted therapies such as BRAF/MEK inhibition and immunotherapies like anti-PD1 agents have shown promising outcomes, an underappreciated possibility is that these two modalities do not work synergistically together but instead undermine each other. Specifically, we propose that BRAF/MEK inhibition unexpectedly, and counterproductively, induces a tumor microenvironment resistant to immunotherapy^1,2^. Remaining unexplored is how melanoma cells reprogram intrinsic signaling pathways after BRAF/MEK inhibition leading to both acquisition of targeted therapy resistance and evasion of immune surveillance.

To understand how this reprogramming might erode antitumor immunity, it is first necessary to consider the immune barriers that melanoma cells must overcome to evade immune surveillance. Multiple groups have established mechanisms through which melanoma tumor cells escape immune surveillance^3,4^. From this body of work, two well-established obstacles have emerged leading to dampened antitumor immunity: one is the minimal intratumoral infiltration of immune cells, termed as immune “coldness”, and the other is the suppression of optimal T cell activation^5–7^. It is unknown whether resistance mechanisms activated by targeted therapy actively contribute to either of the obstacles above, or if their co-occurrence is mechanistically unrelated. Indeed, BRAF/MEK inhibition may carry a hidden cost through its off-target inhibition of anti-tumor immunity.

Among the immune cells that shape whether the tumor microenvironment is permissive or hostile to an antitumor response, macrophages occupy a particularly pivotal position. Macrophages play dual roles in antitumor immunity due to their plasticity^7^. Generally, macrophages can be M1 macrophages or M2 macrophages, exhibiting either anti-tumor or pro-tumor phenotype, respectively leading to either an immune-active or immune-suppressive microenvironment^8,9^. We still lack critical understanding of how melanoma cells dictate the M1/M2 phenotypes of macrophages.

Macrophage polarization state is in turn a critical determinant of whether T cells, the primary effectors of antitumor adaptive immunity, can successfully infiltrate and operate within the tumor. T cells are the key mediators of adaptive immunity against malignancies^10–12^. Unfortunately, most melanomas exhibit minimal intratumoral infiltration of T cells. Even in the case of sufficient T cell infiltration, M2-like macrophages frequently suppress activation of T cells, limiting their functionality^13–15^. Many efforts have subsequently focused on mechanisms to promote intratumoral infiltration of T cells to elicit antitumor immunity.

Beyond the contributions of individual immune cell types, the spatial and functional organization of the immune response within the tumor, exemplified by Tertiary Lymphoid Structures (TLSs), represents an additional and underexplored dimension of antitumor immunity. TLSs function as ectopic lymphoid organs that develop in non-lymphoid tissues typically at sites of chronic inflammation or within tumors^16,17^. Generally, they are comprised of a T cell-rich zone with mature dendritic cells (DCs) adjacent to a B cell follicle, along with high endothelial venules^16–19^. As an immune structure, TLSs mimic the function of lymph nodes. Interestingly, the presence of TLSs in tumor lesions positively correlates with better prognosis^20,21^; however, it remains poorly understood what initiates formation of these immune structures, and importantly how melanoma cells counter immune responses in these structures to evade immunosurveillance. We propose that modeling TLS structures can significantly advance the understanding of antitumor immunity and provide new strategies for immunotherapies.

A molecular entry point into all of these processes, macrophage polarization, T cell infiltration, and TLSs formation, may lie in the transcriptional programs that melanoma cells inherit from their developmental cell of origin. Immature melanocytes such as pigment cells and trophoblast cells highly express Transcription Factor AP-2 Alpha, also known as TFAP2A or AP-2a (Protein Atlas dataset). Frequently misunderstood, TFAP2A or AP-2a is not a family member of AP-1 which is composed of c-Jun and c-Fos. Rather, TFAP2A belongs to the AP-2 transcription factor family, which includes five members: TFAP2A, TFAP2B, TFAP2C, TFAP2D, and TFAP2E^22–24^. *In vivo* mouse studies show TFAP2A is essential for cranial closure and craniofacial development and homozygous TFAP2A knockout results in embryonic lethality^22^.

Several studies suggest that TFAP2A is involved in neural crest development in zebrafish ^23–25^. Notably, melanocytes arise from the neural crest, a transient embryonal structure whose developmental regulatory pathways including TFPA2A are frequently reactivated during tumor progression and phenotypic plasticity. Such plastic states are often implicated in conferring drug resistance. This developmental reactivation is clinically significant because melanoma cells undergoing phenotypic switching to a proliferative and invasive state often co-opt normal developmental neural pathways to evade targeted therapies, suggesting that TFAP2A may act as a node linking embryonic identity to acquired resistance.

Building on this developmental rationale, we investigated whether TFAP2A is induced by BRAF/MEK inhibition and whether it mechanistically connects drug resistance to the immune evasion barriers described above. Herein, we demonstrate that TFAP2A modulates both drug resistance and antitumor immunity, priming melanoma cells to proliferate unchecked. Upregulated TFAP2A confers melanoma cells with drug resistance to BRAF/MEK inhibitors while simultaneously suppressing antitumor immunity. We found that in an immune-compromised mouse model, TFAP2A-KO melanomas show enhanced macrophage infiltration. In a syngeneic mouse model, TFAP2A depletion triggers formation of TLSs. Single cell transcriptomics further confirms a favorable microenvironment for antitumor immunity in the absence of Tfap2a, with more anti-tumor M1-like macrophages, less pro-tumor M2-like macrophages, more intratumoral T cells and more mature dendritic cells. Finally, clinical datasets reveal a significant correlation of TFAP2A expression with prognosis and immune signatures, underscoring TFAP2A as a convergent regulator of drug resistance and anti-tumor immunity.

## Results

### BRAF inhibition upregulates a discrete set of transcription factors with TFAP2A emerging as the top candidate linked to secondary addiction

BRAF^V600E^ is the dominant oncogenic driver in melanoma and the primary target of frontline targeted therapy^26–29^; however, melanoma invariably develops resistance under sustained BRAF/MEK inhibition, indicating that tumor cells can rewire their transcriptional dependencies, from BRAF addiction to alterative survival pathways.

We hypothesized that factors upregulated specifically in response to sustained BRAF/MEK inhibition represent candidate drives of “secondary addition.” To profile global transcriptomic changes induced by BRAF suppression, we used two orthogonal approaches including pharmacological inhibition with Dabrafenib/Trametinib (Dab/Tra), and CRISPR/CAs9 mediated BRAF knockout in a patient derived xenograft (PDX) model WND238. This model undergoes dramatic initial tumor regression in response to Dab/Tra treatment followed by a static stage or minimal residual diseases for about 6 months, with later relapse (**Fig 1a**).

**Figure 1.**
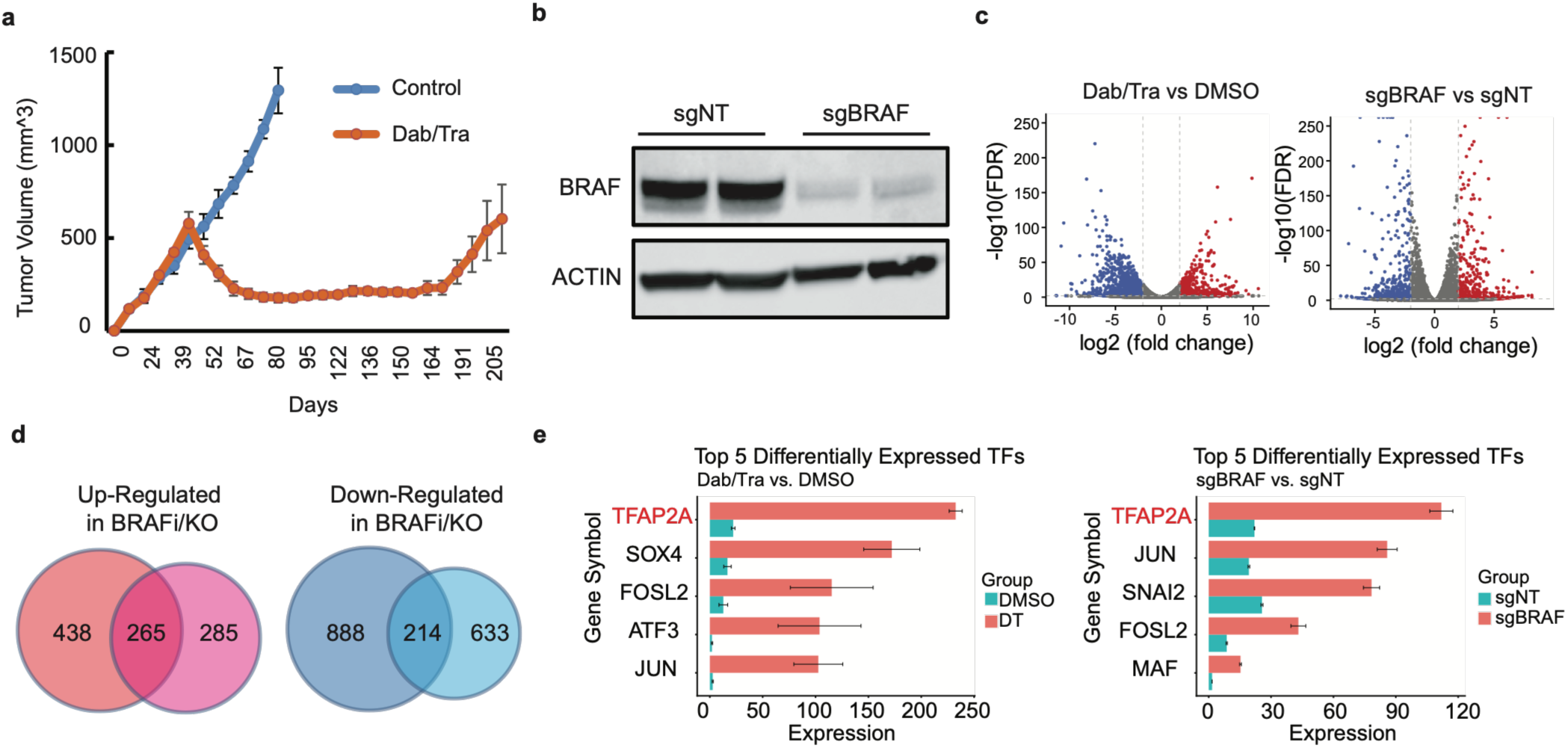
Identification of TFAP2A as top candidate of transcription factor involved in drug resistance. **(a)** WND-238 human melanoma cells were subcutaneously injected into NSG mice for tumor growth curve. Melanomas were treated with or without BRAF/MEK inhibitors, Dabrafenib/Trametinib (Dab/Tra), when they reach around 500mm^3^. **(b)** A single guide RNA targeting BRAF was designed and cloned to LentiV2 backbone for virus production. WND238 cells were infected with non-targeting sgRNA (sgNT) or BRAF-targeting sgRNA (sgBRAF) and selected with blasticidin. Western blot to confirm the knockdown efficiency in the pooled sgBRAF compared to sgNT. **(c)** Bulk-RNA sequencing to identify the differentially expressed genes (DEGs) by comparing WND238 cells treated with Dab/Tra to the DMSO control **(left panel)** or WND238 cells infected with sgBRAF to the sgNT control **(right panel)**. Volcano plot indicates the upregulated (red color) or downregulated DEGs (blue color). **(d)** Overlapped upregulated genes or downregulated genes by comparing the DEGs from pharmacological inhibition of BRAF **(c, left panel)** and genetic disruption of BRAF (c, right panel). 265 upregulated genes and 214 downregulated genes were overlapped, which is regarded as BRAF-*regulatome*. **(e)** From the 265 upregulated genes regulated by BRAF, we further narrowed down to the top five transcription factors and TFAP2A is the top one candidate.

We next established BRAF knockout cells mediated by CRISPR/Cas9 genome editing and confirmed that BRAF protein level is significantly downregulated compared to non-targeting control (sgNT) (**Fig 1b**). Next, we performed bulk RNA-seq across both perturbation conditions to identify the transcriptomic changes by comparing Dabr/Tra treated cell and BRAF knockout cells with their respective controls (**Fig 1c**). To rule out off-target effects from either inhibitor or CRISPR/Cas9, we overlapped the up- or down- regulated genes shared by Dra/Tra treatment and BRAF knockout cells, which are more reliably BRAF-regulated genes, and termed them “*BRAF-regulatome*” (**Fig 1d**).

Among the upregulated genes, we decided to focus on transcription factors (TFs) reasoning that they regulated entire downstream gene networks and are more likely positioned to reprogram cell states and confer resistance. TFAP2A ranked as our top hit and was the least characterized in the resistance context, making it our top candidate for further investigation (**Fig 1e**). Other top hits including JUN, FOSL2, SOX4, SNAI2 were relatively well characterized in melanoma resistance, and therefore not pursued herein^30–32^.

### BRAF/MEK inhibition drives robust and consistent upregulation of TFAP2A across multiple models

As discussed above, several previous studies suggest resistant melanomas retain neural-crest stem cell-like features, and that TFAP2A contributes to neural crest development^33,34^. We hypothesized that upregulation of TFAP2A following BRAF/MEK inhibition may represent an alternative pathway through which melanoma cells maintain stemness and evade cell death.

Having identified TFAP2A as the top transcription factor in the BRAF regulatome, we next validated its upregulation by BRAF/MEK inhibition across independent datasets and experimental systems. Our bulk RNA sequencing confirmed that TFAP2A mRNA was significantly elevated in response to both Dab/Tra treatment and sgRNA mediated BRAF inhibition compared to control (**Fig 2A)**. Western blot analysis secondarily confirmed Dab/Tra treatment incudesTFAP2A protein (**Fig 2b**).

**Figure 2.**
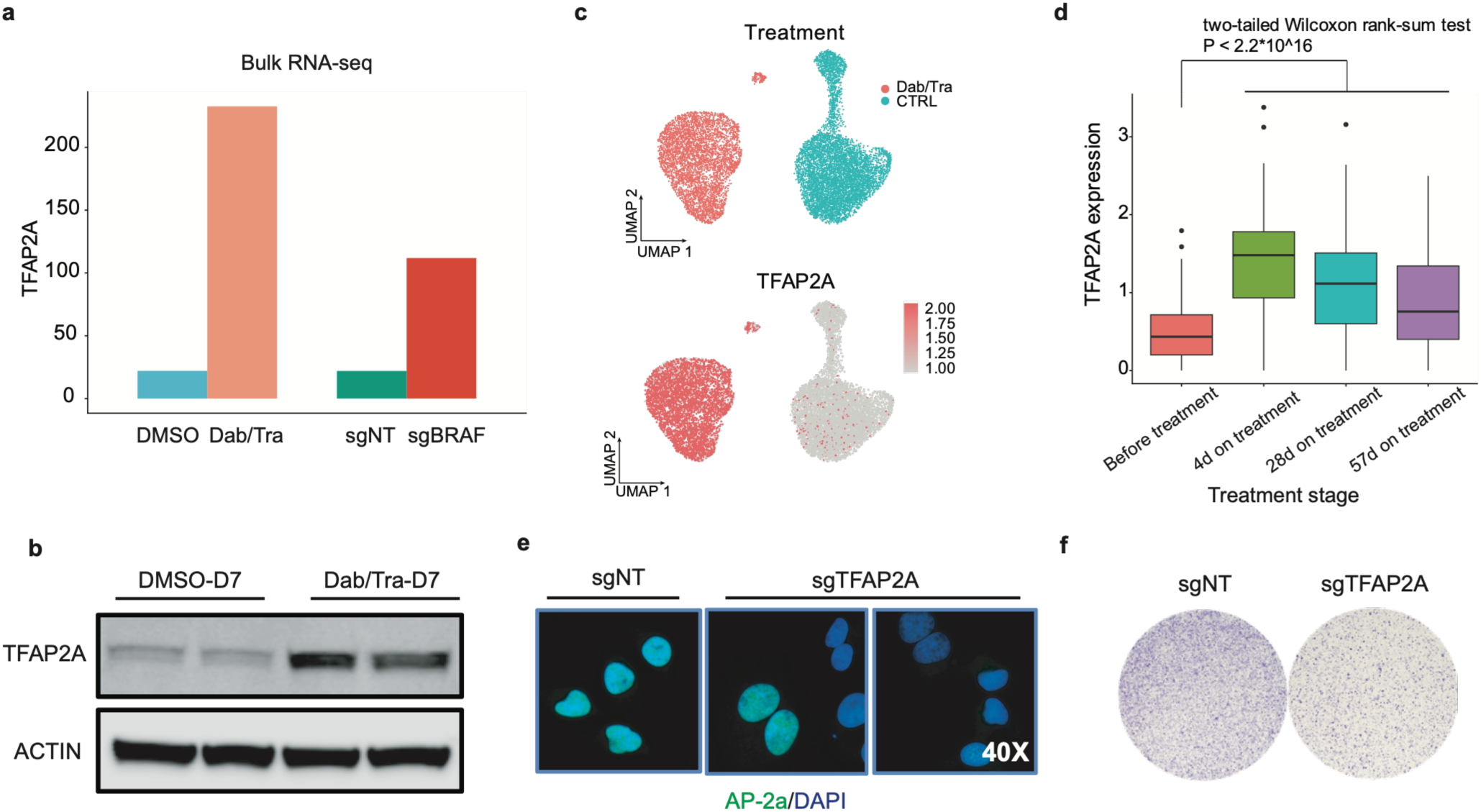
TFAP2A is upregulated by BRAF/MEK inhibition, with its genetic disruption impairing cell proliferation *in vitro*. (**a**) WND238 cells were treated with Dab/Tra and DMSO control, or infected with sgRNAs targeting BRAF (sgBRAF) and sgNT control, respectively. Total mRNAs were isolated for bulk RNA-seq. TFAP2A mRNA level was normalized and presented as indicated. (**b**) WND238 cells were trated with Dab/Tra or DMSO. Total protein lysates were collected for western blot to detect the TFAP2A protein expression. (**c**) Single cell RNA sequencing (scRNA-seq) from published dataset to detect the TFAP2A mRNA expression by comparing the Dab/Tra treated melanomas and their controls. Upper panel shows the treatment condition while the lower panel shows TFAP2A is upregulated in the Dab/Tra treated condition. (**d**) Another independent single cell dataset confirms that TFAP2A is upregulated by BRAF/MEK inhibitors, Dab/Tra. (**e**) WM4380-2 cells, which are BRAF-MEK-inhibition resistant, were infected with sgRNA targeting TFAP2A (sgTFAP2A) or non-targeting control (sgNT). Immunofluorescent staining was used to detect TFAP2A protein. (**f**) The same treatment as (e). 10,000 cells were seeded onto 6-well plate and cultured for 10 days for colony formation. Crystal violet solution was used to visualize the colonies.

To extend these findings beyond our own experimental models, we utilized published single cell RNA-sequencing (scRNA-seq) data^35^ and compared the TFAP2A mRNA expression of melanoma cells treated with Dab/Tra with that of control cells. Consistently, uniform manifold approximation and projection (UMAP) showed that the cell cluster treated with Dab/Tra had significantly higher expression of TFAP2A (**Fig 2c**). A second independent public dataset^33^ also supported our findings that BRAF/MEK inhibition upregulates TFAP2A (**Fig 2d**). Taken together, these results establish TFAP2A upregulation as a reproducible and consistent transcriptional response to BRAF/MEK inhibition, consistent with a role of promoting melanoma cell survival under treatment pressure.

### TFAP2A disruption reduces proliferation of melanoma cells *in vitro*

To test whether and to what extent survival of BRAF/MEK-inhibition resistant melanoma cells are dependent on TFAP2A, we established TFAP2A knockout cells using CRISPR/Cas9 genome editing in a BRAF/MEK-inhibition resistant model, WM4380-2. Notably, this PDX model most closely resembles primary resistance, as this patient did not respond to BRAF/MEK inhibition at the outset. We infected WM4380-2 cells with lentiviruses carrying non-targeting sgRNA (sgNT) or TFAP2A-targeting sgRNA (sgTFAP2A) and pooled all surviving cells selected with blasticidin. We then performed immunofluorescence (IF) to detect TFAP2A protein levels and confirmed that the sgTFAP2A pooled cells show robust knockout phenotype compared to sgNT cells (**Fig 2e**). Colony formation assays showed sgTFAP2A treated cells have impaired growth compared to sgNT cells (**Fig 2f**).

### Genetic disruption of TFAP2A overcomes BRAF/MEK inhibitor resistance *in vivo,* inducing a stromal remodeling phenotype

Given that CRISPR/Cas9 mediated pooled TFAP2A knockout cells are a mix of genotypes, we picked single clones of TFAP2A knockout (TFAP2A-KO) cells and confirmed TFAP2A protein loss by western blot and sequencing (**Fig 3a, 3b**). We then subcutaneously injected the TFAP2A-KO cells or their wildtype control (TFAP2A-WT) cells into NSG mice to compare tumor growth in the BRAF/MEK- inhibition resistant model, WM4380-2. As anticipated, TFAP2A-KO melanomas show significantly decreased tumor growth compared to wildtype controls (**Fig 3c**). *In situ* hybridization (ISH) confirmed the absence of TFAP2A mRNA in knockout tumors (**Fig 3d**).

**Figure 3.**
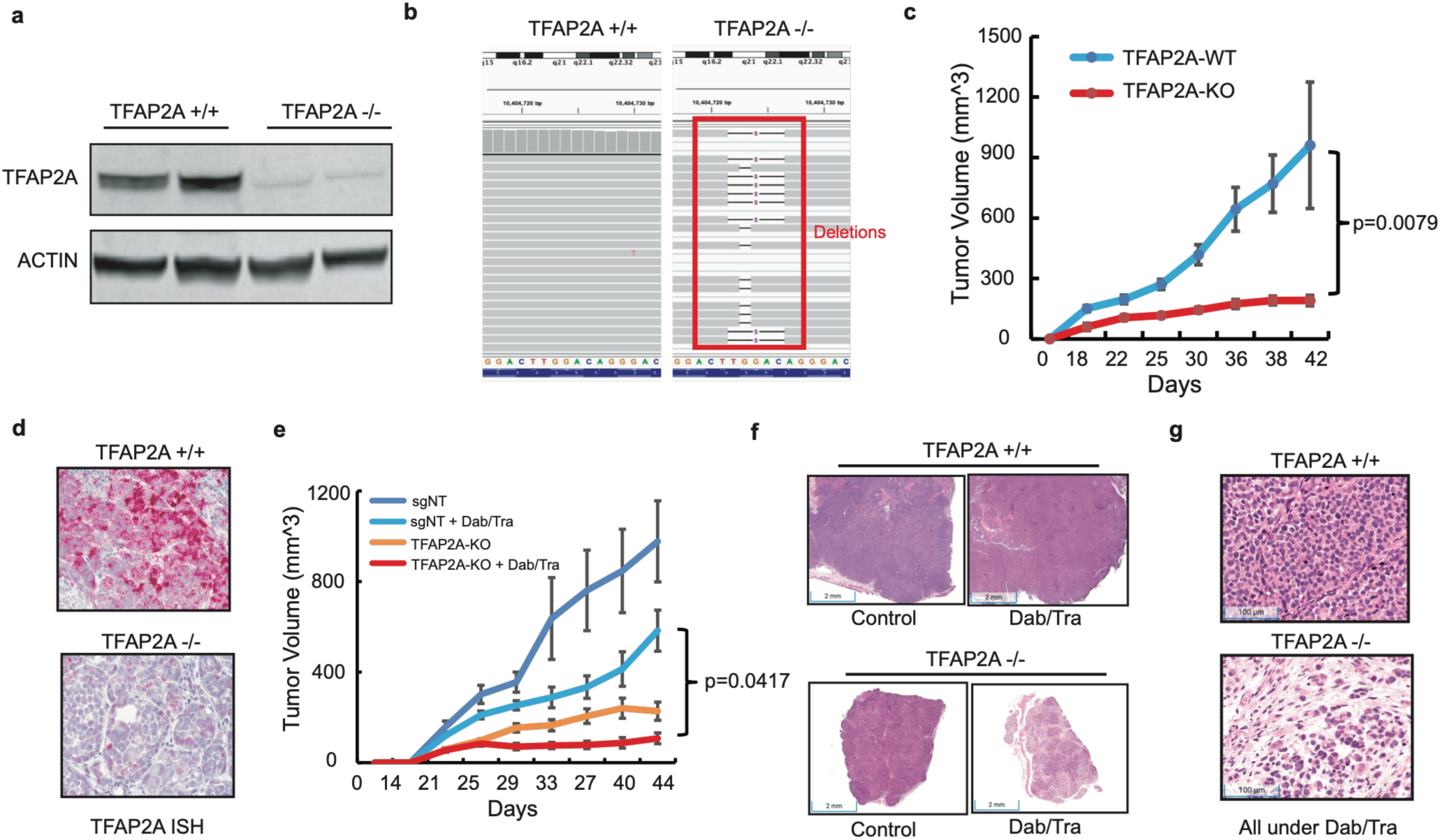
TFAP2A-KO melanomas overcome BRAF/MEK-inhibition resistance and exhibit distinct histology with more stromal enrichment. (**a**) WM4380-2 cells were infected with lentiviruses carrying sgRNA targeting TFAP2A and selected with Blasticidin. Single colonies were picked by and expanded. Western blot was used to validate the loss of TFAP2A protein in the single clone. (**b**) Single TFAP2A-KO cells or the wild-type cells were subjected to bulk RNA-seq for detection of small insertion or deletions (indels). Single TFAP2A-KO cells show small deletions resulting in frameshift while the wildtype do not have indels. Red arrows indicate the small deletions. (**c**) TFAP2A-WT or TFAP2A-KO melanoma cells were subcutaneously injected into NSG mice for tumor growth curve (n=5). (**d**) Tumors from the same treated in (c) were collected for histology. *In situ* hybridization was used to detect the mRNA expression of TFAP2A. (**e**) TFAP2A-WT or TFAP2A-KO cells were subcutaneously injected into NSG mouse for tumor growth. Mice were then treated with Dab/Tra chow or control diet around day 10 as indicated in the figure (n=5). (**f** and **g**) The same treatment from (e), tumors were collected for H&E staining. TFAP2A-KO melanomas with Dab/Tra treatment exhibit distinct histology compared to all other groups, with more stromal enrichment. Higher magnification (g) indicates between the melanoma islands, there are more extracellular matrix and stromal cells.

Given that TFAP2A is upregulated by BRAF/MEK inhibition (**Fig 2**), and TFAP2A contributes to survival of resistant melanoma cells (**Fig 3c**), we further asked whether TFAP2A disruption overcomes resistance under the drug pressure of BRAF/MEK inhibitors (Dab/Tra) *in vivo*. To test this, we subcutaneously injected TFAP2A-WT and single TFAP2A-KO cells into NSG mice and monitored tumor growth. When tumors were palpable, we treated WT or KO tumor with or without Dab/Tra. Strikingly, we observed TFAP2A-KO melanomas treated with Dab/Tra maintained the size for more than three weeks and showed significantly decreased growth compared with that of TFAP2A-WT melanomas (**Fig 3e**), suggesting in the context of BRAF/MEK inhibition, TFAP2A is essential for tumor growth. Intriguingly, H&E staining revealed that only TFAP2A-KO melanomas treated with Dab/Tra show a distinct histology compared to all other groups (**Fig 3f and 3g**), with more eosin positive space between cell nuclei and spindle-like cells between melanoma islands, indicating more stromal enrichment. Interestingly, we also found that TFAP2A-KO tumors show less pigmentation. We hypothesized this might be due to less Microphthalmia-Associated Transcription Factor (MITF) expression in the absence of TFAP2A (**Sup Fig 1**).

### TFAP2A crosstalk with BRAF/MEK pathway reprograms the transcriptome and promotes intratumoral macrophage infiltration

Given that TFAP2A expression is induced by BRAF/MEK inhibitors (**Fig 2**) and genetic disruption of TFAP2A overcomes BRAF/MEK-inhibition resistance (**Fig 3e**), we hypothesized TFAP2A acts as a transcriptional relay within the BRAF/MEK signaling axis, reprogramming gene expression to support cell survival under drug pressure. To test this, we performed bulk RNA-seq to identify the differentially expressed genes (DEGs) from TFAP2A wildtype or knockout melanomas treated with or without Dab/Tra as indicated in **Fig 4a**.

**Figure 4.**
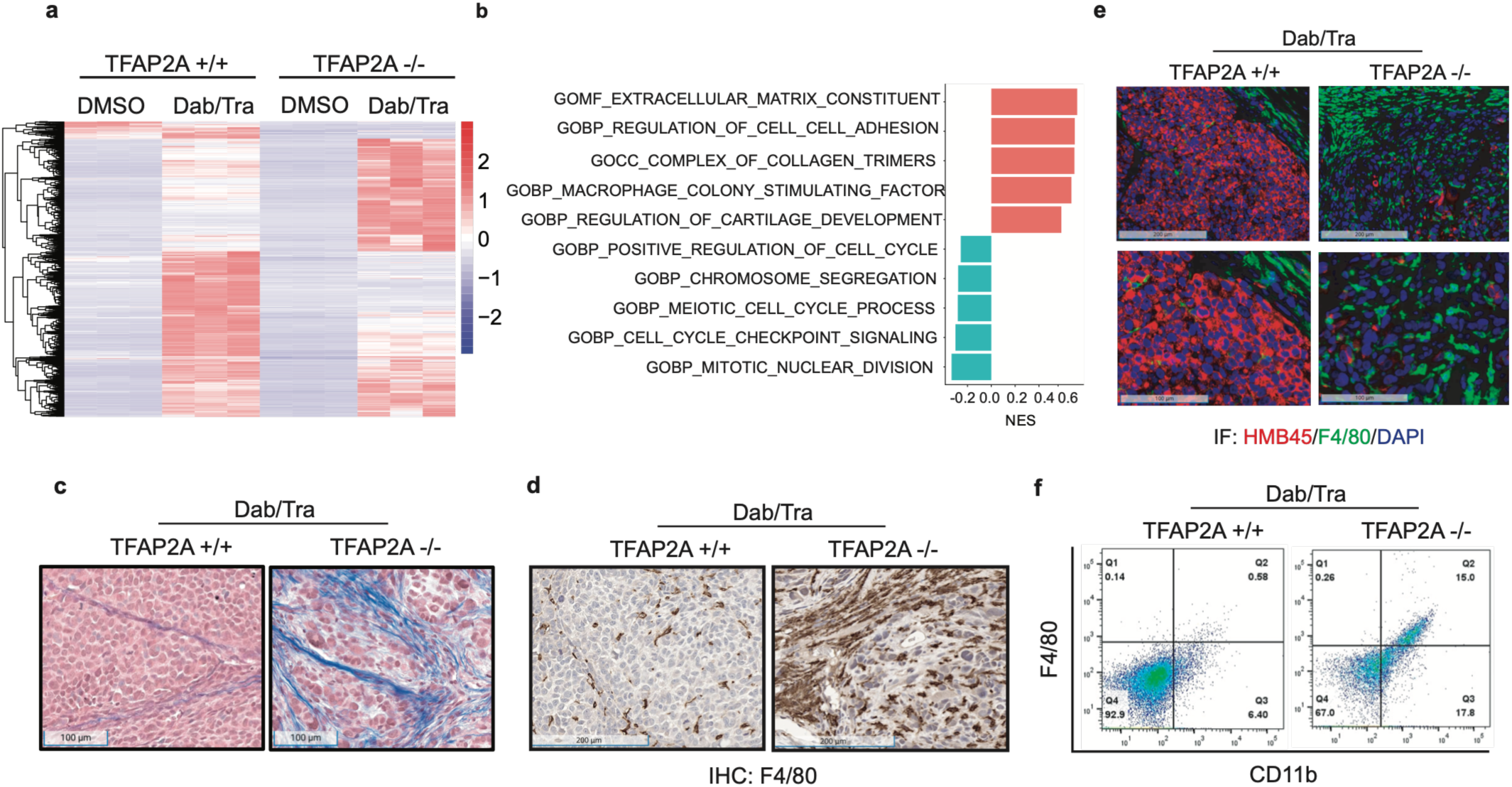
TFAP2A-KO melanomas exhibit enriched collagen and enhanced infiltration of macrophages. (**a**) WM4380-2 TFAP2A-WT or TFAP2A-KO melanoma cells were treated with DMSO or Dab/Tra. Total mRNAs were isolated for bulk RNA-seq. Differentially expressed genes (DEGs) were used to draw the heatmap. TFAP2A-KO cells treated with Dab/Tra reveal a unique gene signature compared to other groups. (**b**) GSEA analysis using DEGs from (a) shows extracellular matrix is enriched in TFAP2A-KO cells treated with Dab/Tra. (**c**) Trichrome staining shows more collagen positive signal (blue color) in TFAP2A-KO melanoma treated with Dab/Tra. (**d**) IHC staining shows more F4/80+ macrophages detected in TFAP2A-KO melanoma treated with Dab/Tra. (**e**) Immunofluorescent co-staining of HMB45 and F4/80 confirms that TFAP2A-KO melanoma expresses less HMB45, while with more intratumoral infiltrated F4/80+ macrophages. (**f**) FACS analysis to quantify the F4/80+Cd11b+ macrophages in TFAP2A-KO melanoma treated with Dab/Tra compared to its control.

Intriguingly, Heatmap from DEGs revealed a unique gene signature in the group of TFAP2A-KO cells treated with Dab/Tra compared to all other groups (**Fig 4a**), indicating crosstalk betweenTFAP2A and BRAF/MEK pathways. Focusing on the comparison between TFAP2A-WT and TFAPP2A-KO both under treatment pressure, Gene Sets Enrichment Analyses (GSEA) identified enrichment of cell matrix pathways in TFAP2A-KO melanomas treated with Dab/Tra (**Fig 4b**). This finding was further validated with trichrome staining and confirmed the enrichment of collagens in TFAP2A-KO melanomas treated with Dab/Tra (**Fig 4c**).

Importantly, the GSEA analyses also revealed an immune signature in the TFAP2A-KO melanomas treated with Dab/Tra. Immunohistochemistry (IHC) staining demonstrated F4/80+ macrophages were significantly more abundant as infiltrate into the TFAP2A-KO melanomas treated with Dab/Tra compared to the wildtype treated with Dab/Tra (**Fig 4d**). Immunofluorescent co-staining of melanoma marker HMB45 and macrophage marker F4/80 corroborated enhanced intratumoral macrophage infiltration in TFAP2A-KO (**Fig 4e**). Moreover, flow cytometry reached the same conclusion that TFAP2A-KO melanomas treated with Dab/Tra contain more CD11b+ and F4/80+ macrophages (**Fig 4f**) than the wildtype controls. ChIP-seq further identified the direct transcriptional targets of TFAP2A by comparing pulled down DNA fragments from TFAP2A-WT and TFAP2A-KO cells (**Sup Fig 2**). GSEA analysis demonstrated that most of the TFAP2A targets were in neurogenesis and axonogenesis pathways (**Sup Fig 2**).

Taken together, these data demonstrate that TFAP2A orchestrates a transcriptional program downstream of BRAF/MEK signaling that suppresses stromal remodeling and immune infiltration, and further that disruption of TFAP2A restores both processes.

### Tfap2a depletion triggers Tertiary Lymphoid Structures (TLSs) and potent immunity in an immunocompetent model

To this point, our work had been conducted in NSG immunocompromised mice, which lack functional lymphocytes and are therefore unsuitable for further study of adaptive immunity. To better characterize the role of TFAP2A in anti-tumor immunity, we next switched to a syngeneic model using murine melanoma cell line YUMM1.7 in immune-competent C57BL/6J mice.

First, we employed CRISPR/Cas9 gene editing to establish the Tfap2a-KO cells and confirmed loss of Tfap2a in single knockout clones (**Fig 5a**). Strikingly, while wildtype tumors grew progressively, Tfap2a-KO tumors formed small nodules around day 10 and then regressed robustly (**Fig 5b**). This phenotype was independently replicated in the B16F10 murine melanoma model (**Sup Fig 3**). Histological examination of TFAP2A KO tumors revealed a distinctive architecture similar to what was observed in the NSG mouse model. Specifically, we observed, expanded eosinophilic space in the tumor center surrounded by a dense cluster of cells with condensed, small nuclei consistent with organized immune infiltrates (**Fig 5c**). Immunofluorescence (IF) staining confirmed co-localization of Cd8+ T cells with aggregates of Cd20+ B cells, a hallmark of tertiary lymphoid structures (TLSs) (**Fig 5d**). Confocal immunofluorescent staining additionally revealed CD11c+ dendritic cells within these immune aggregates, completing the canonical TLSs (**Fig 5e**).

**Figure 5.**
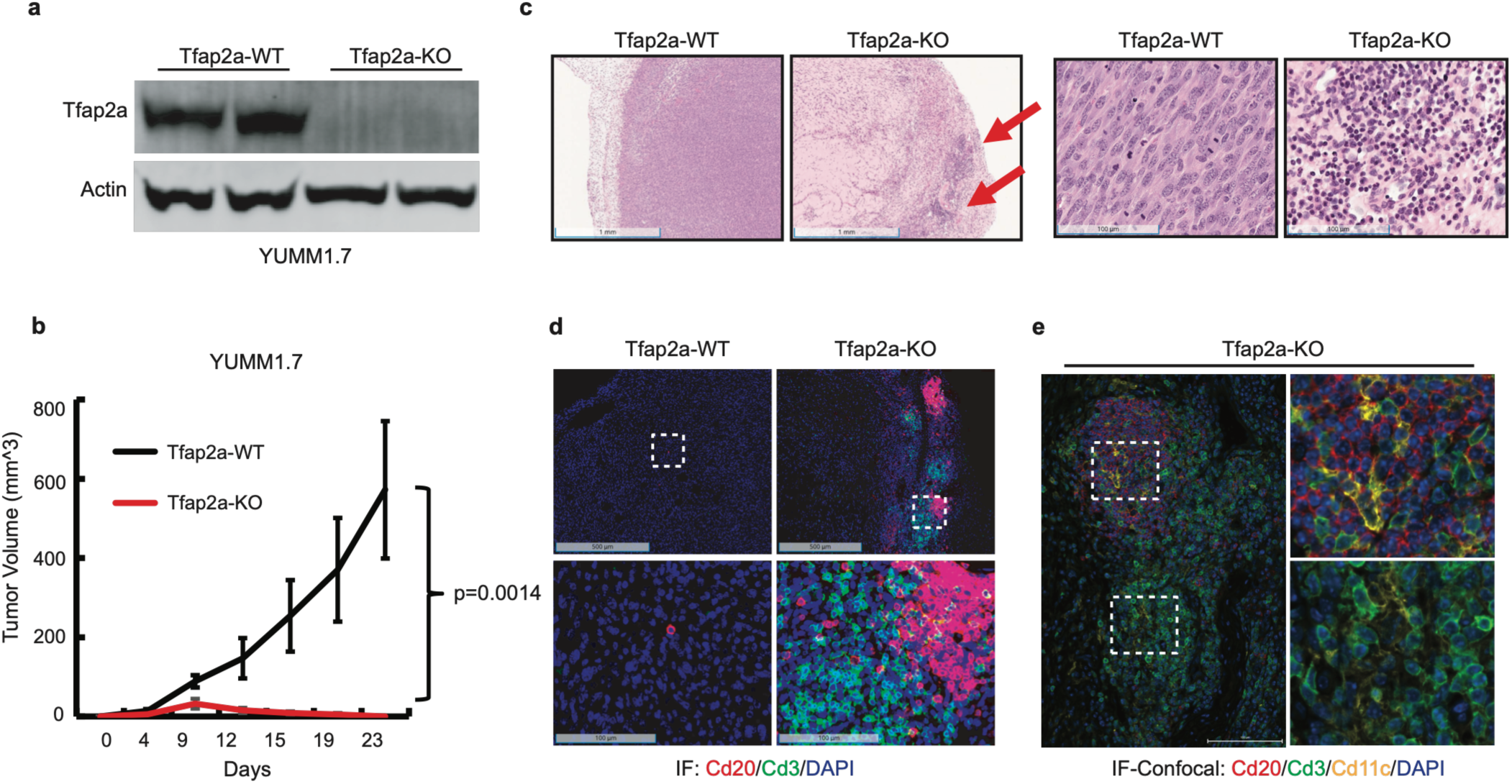
Genetic disruption of Tfap2a impairs tumor growth and induces tertiary lymphoid structures (TLSs) in an immune-competent mouse model. (**a**) YUMM1.7 cells were infected with lentiviruses carrying CRISPR/Cas9 cassette to targeting mouse Tfap2a. Single knockout cells were picked up, expanded and validated by western blot to confirm the loss of Tfap2a protein. Beta-actin was used as a loading control. (**b**) YUMM1.7 Tfap2a wildtype (Tfap2a-WT) or knockout (Tfap2a-KO) cells were subcutaneously injected into C57BL/6J mice for tumor growth curve (n=5). (**c**) Tfap2a-KO melanoma shows a distinct histology with more matrix-enriched area and cell cluster with small nuclei, indicating immune cells. (**d**) Immunofluorescent co-staining of Cd20, Cd3 shows T cell clusters and B cell clusters which are adjacent to each other, resembling tertiary lymphoid structure. (**e**) Confocal immunofluorescent staining shows Cd11c+ dendritic cells within the TLSs.

Interestingly, we found Yumm1.7 cells are responsive to BRAF/MEK inhibitors, showing dramatic tumor shrinkage under the drug pressure. However, tumor cells quickly acquire resistant mechanisms and relapsed (**Sup Fig 4**). Most interestingly, we observed a significant intratumoral infiltration of Cd3+ T cells in the acute phase of BRAF/MEKi treatment, while in the relapsed melanomas, Cd3+ T cells are expelled from tumors and become an immune-cold feature again (**Sup Fig 4**). This suggests our mouse model not only mimic the tumor growth but also the immune signature in melanoma patients, especially resemble the immune-cold signature of relapsed tumors. Of note, Yumm1.7 is also an anti-PD1 resistant model.

Taken together, we demonstrate that Tfap2a depletion alone is sufficient to initiate the assembly of an organized anti-tumor immune structure TLSs within the tumor in an immunotherapy resistant model.

### Single cell transcriptomics reveal Tfap2a loss comprehensively remodels the tumor immune microenvironment toward an anti-tumor state

To further dissect the tumor microenvironment reshaped in the absence of Tfap2a, we performed single cell RNA sequencing (scRNA-seq) to compare the immune components of Tfap2a-KO melanoma to the wildtype control (**Fig 6a**). Bioinformatic analysis identified the differentially expressed genes and annotated different types of cells including melanoma cells, immune cells, fibroblasts and endothelial cells (**Fig 6b, Sup Fig 5**). Consistent with our histological findings, in TFAP2a-KO tumors we observed markedly increased intratumoral infiltration of T cells and B cells (**Fig 5d, 6c and 6e**). Macrophage phenotyping revealed a striking reprogramming of macrophage plasticity. Tfap2a-KO tumors contained more M1-like antitumor macrophages and fewer M2-like immunosuppressive macrophages, indicating a shift toward an immune-active microenvironment (**Fig 6c and 6d**). Further, scRNA-seq analyses also revealed an increase of matured dendritic cells (DCs), the professional antigen presenting cells (APCs), in the immune component of the Tfap2a-KO melanoma (**Fig 6f**), consistent with the staining results (**Fig 5e**). Cell-cell communication analyses further showed that TGF-beta signaling network is reduced in the Tfap2a-KO melanoma (**Fig 6g**), further supporting a less immunosuppressive microenvironment.

**Figure 6.**
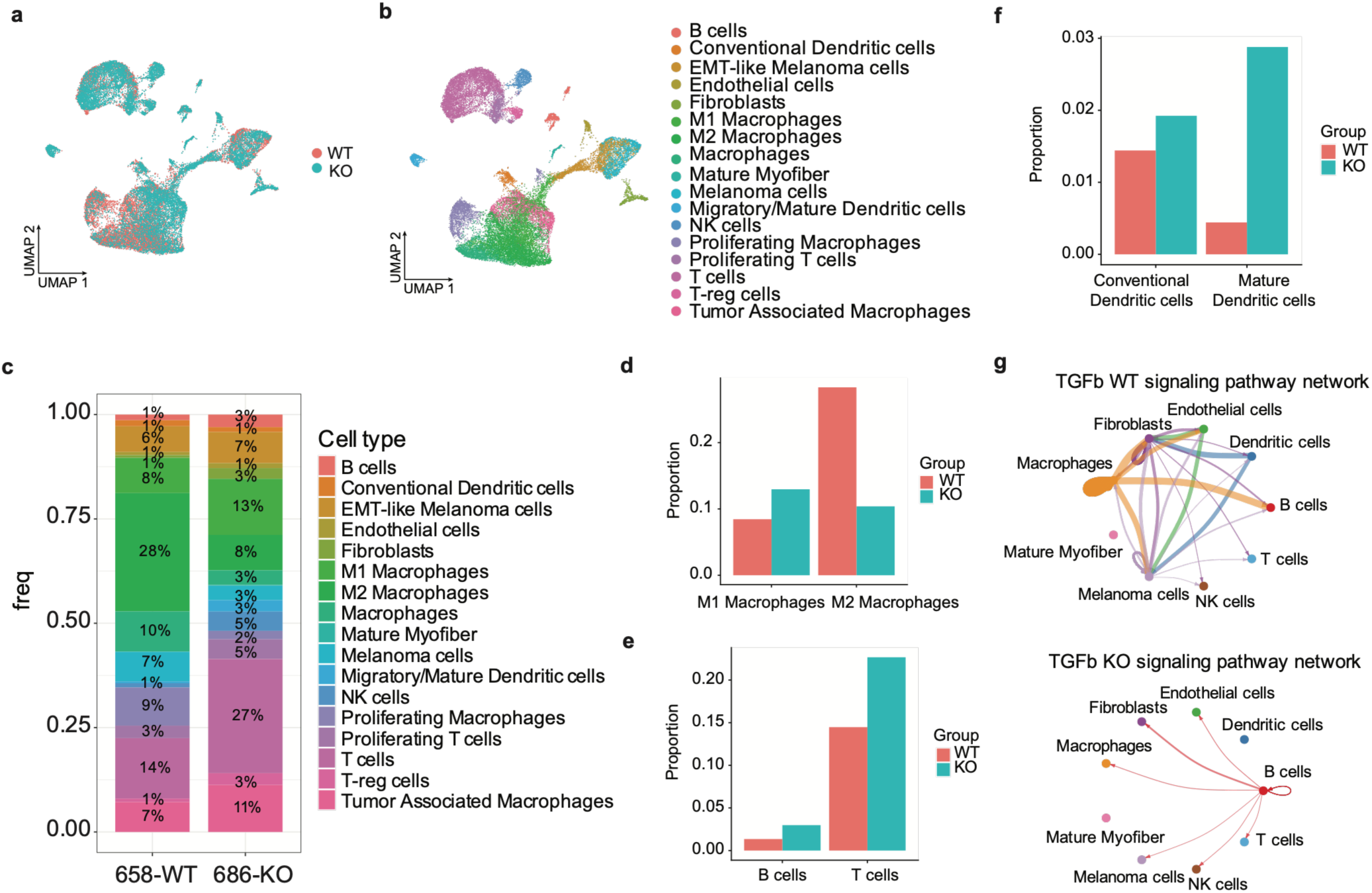
Single cell transcriptomics profile the landscape of tumor microenvironment within Tfap2a-KO melanoma, supporting a less immunosuppressive environment with more M1-like and less M2-like macrophages. (**a** and **b**) Single cell RNA sequencing analysis of YUMM1.7 Tfap2a-WT or Tfap2a-KO melanomas to profile the cell components of the tumor microenvironment. Differentially expressed genes were used for annotation of cell types. (**c**) Individual cell percentage of annotated cell types in (b). (**d**) Tfap2a-KO melanoma shows more M1-like and less M2-like macrophages. (**e**) Tfap2a-KO melanoma reveals more B cells, NK cells, and T cells while less T-regs within the microenvironment. (**f**) More mature DCs were observed in the Tfap2a-KO melanoma compared to the wildtype control. (**g**) Cell-Cell-Communication (CCC) reveals much less cell-cell interactions of TGF-beta signaling in the Tfap2a-KO melanoma.

Collectively, these single cell data reveal that TFAP2a loss does not simply augment one arm of the immune response but comprehensively restructures the tumor immune microenvironment promoting antitumor macrophage polarization, T and B cell infiltration, dendritic cell maturation and suppression of immunosuppressive cytokine signaling.

### High expression of TFAP2A in melanoma patients correlates with poor prognosis and suppressed immune gene signatures

To ask whether our findings have clinical significance, we utilized The Cancer Genome Atlas Program (TCGA) database and performed correlative analyses. We sub-grouped the melanoma patients based on the TFAP2A expression level as low, medium or high expression (**Fig 7a**). We found TFAP2A high expression correlates with significantly worse overall survival (**Fig 7a**). Differential gene expression analysis comparing the high versus low/medium TFAP2A groups showed that genes upregulated in the low/medium TFAP2A group exhibited a graded increase as TFAP2A expression declined (**Fig. 7b**), suggesting a dose-dependent relationship between TFAP2A suppression and a pro-immunity transcriptional program (**Fig 7b**). Furthermore, we used the DEGs to identify pathways enriched in the low/medium TFAP2A group compared to the TFAP2A high group. Strikingly, five immune-related pathways are enriched in the TFAP2A low/medium group including inflammatory responses, IL6-JAK-STAT3, interferon-gamma responses, TNFA signaling and IL2-STAT5 signaling (**Fig 7c**), which supports the conclusion from mouse model that genetic disruption of Tfap2a promotes antitumor immunity (**Fig 5 and 6**).

**Figure 7.**
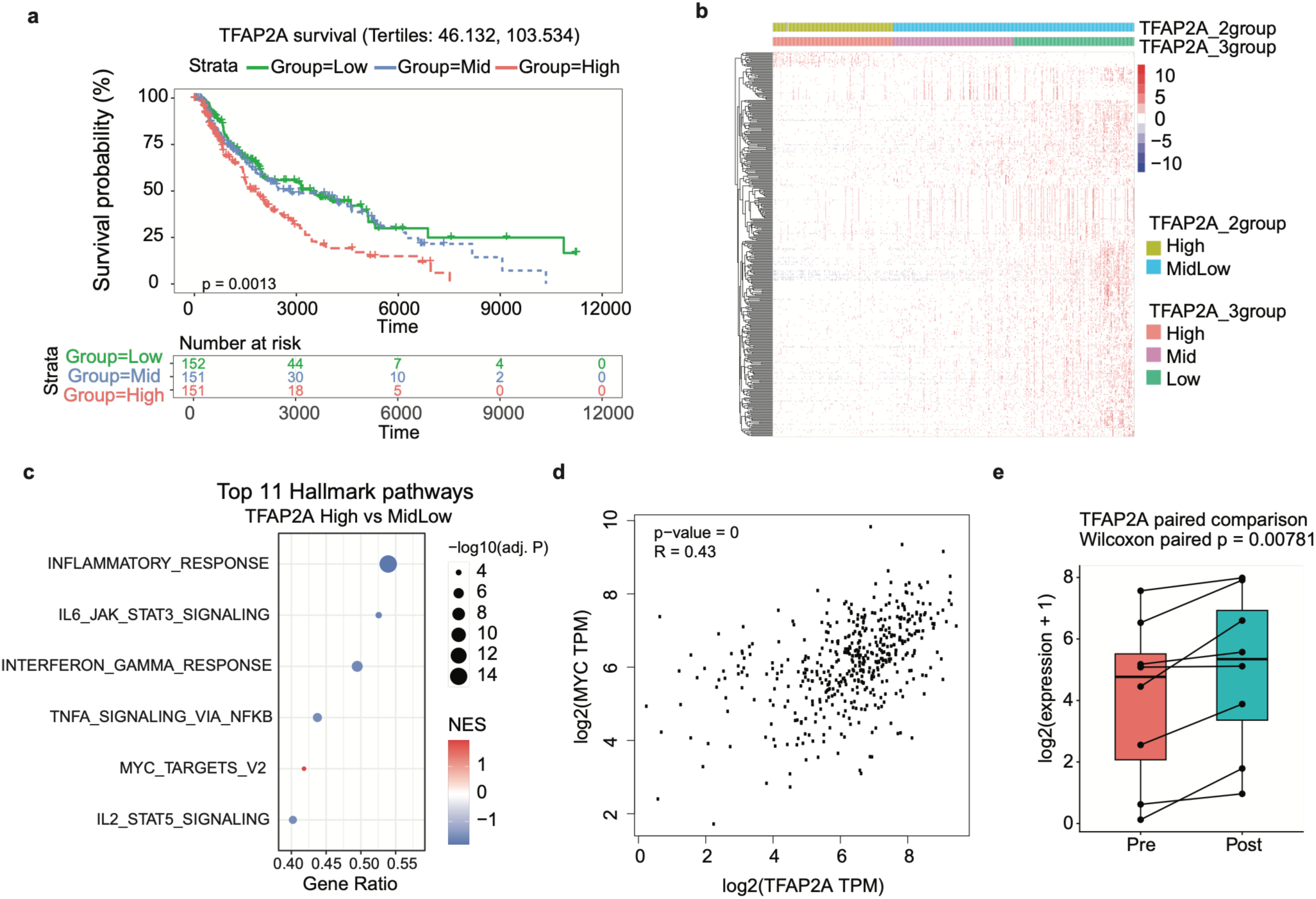
Low expression of TFAP2A correlates with good prognosis and enrichment of immune signatures in melanoma patients. (**a**) Using TCGA dataset, we grouped melanoma patients as TFAP2A low, medium and high expression and performed survival analysis. High TFAP2A expression correlates with worse prognosis, while low/medium correlates with better survival probability. (**b**) Differentially expressed genes (DEGs) by comparing TFAP2A high group with low/medium group. More upregulated DEGS were observed in TFAP2A low/medium group compared to TFAP2A high group. (**c**) GSEA analysis reveals five immune-related pathways are enriched in the TFAP2A low/medium group including inflammatory response, IL6-JAK-STAT3 signaling, interferon-gamma response, TNFA signaling via NF-κB, IL2-STAT5 signaling. MYC targets are more enriched in TFAP2A high group. (**d**) Correlative analysis reveals TFAP2A expression positively correlates with MYC expression in TCGA melanoma patients’ dataset. (**e**) Another public dataset of melanoma PDX models confirm BRAF/MEK inhibitors significantly upregulate TFAP2A expression.

**Figure 8.**
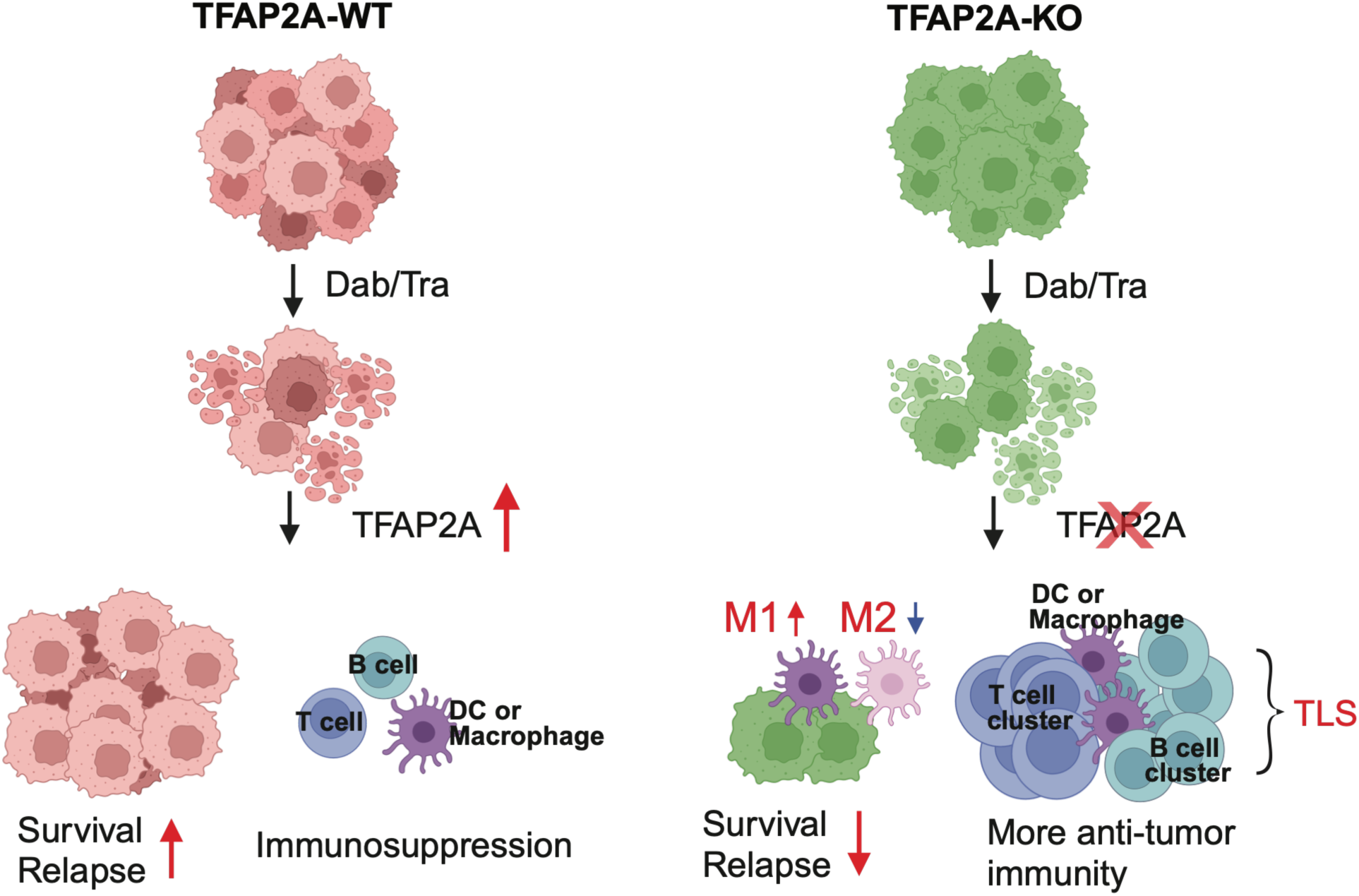
Schematic illustration of the role of TFAP2A in drug resistance and antitumor immunity. In BRAF-mutation driven melanomas, BRAF/MEK inhibitors, Dab/Tra, significantly upregulate TFAP2A which confers melanoma cells resistance and suppresses antitumor immunity, thereby contributing to recurrence. However, in TFAP2A knockout melanomas, the upregulation of TFAP2A is blocked and the conferred resistance is overcome, meanwhile, macrophages are polarized to M1 phenotype, TLSs formation is triggered, thereby antitumor immunity is elicited.

Interestingly, we also observed MYC signaling pathway enrichment in TFAP2A high group (**Fig 7c**). Further analysis confirmed a significant correlation between TFAP2A and MYC expression in TCGA dataset (**Fig 7d**). We next utilized another public dataset, which focused on melanoma PDX models to study BRAF/MEK-inhibition resistance, and found BRAF/MEK inhibitors significantly upregulate TFAP2A (**Fig 7e**), again, consistent with our experimental data (**Fig 2a-2d**). These results from public dataset suggest that TFAP2A is playing a critical role in modulating antitumor immunity and potentially represents a target for therapeutics.

Taken together, these clinical analyses establish that TFAP2A is not only a mechanistic driver of drug resistance and immune response in experimental models, but also a clinically meaningful determinant of immune landscape and patient outcome in melanoma.

## Discussion

Despite the impressive clinical success of BRAF/MEK inhibition in melanoma, acquired resistance remains a near universal obstacle and the molecular pathways that simultaneously drive drug tolerance and suppress anti-tumor immunity are poorly identified. Herein we identify TFAP2A as a transcription factor that is induced by BRAF/MEK inhibition and serves as a convergent driver of both processes. We find that genetic disruption of TFAP2A overcomes inhibitor resistance in part by its ability to reprogram macrophage polarization toward an M1-like anti-tumor state, and formation of tertiary lymphoid structures. Together, this represents a dual therapeutic vulnerability with implications for both targeted therapy and immunotherapy.

### TFAP2A exerts a distinct, context-dependent role in immunocompromised vs immunocompetent settings

Central to interpreting the role of TFAP2A in shaping the melanoma tumor immune microenvironment is careful interpretation of our immunocompromised and immunocompetent *in vivo* data. In the NSG immunocompromised model, which lacks functional T and B cells, TFAP2A knockout tumors treated with BRAF/MEK inhibition showed enhanced intratumoral infiltration of macrophages and stromal remodeling. This model effectively functions to isolate the intrinsic, cell autonomous effects of TFAP2A on the melanocyte and innate immune system recruitment alone due to the lack of adaptive immunity. The macrophage infiltration observed in the NSG model can therefore not be contributed to T or B cell mediated signaling, and we hypothesize instead may rely on cytokine signaling, extra-cellular matrix composition, or even chemokine gradients. All above mechanisms are subject to future investigation.

A significant limitation of this model is that it alone cannot address whether TFAP2A loss is sufficient to drive a complete innate and adaptive anti-tumor response limiting our ability to understand the full functional consequents of macrophage reprogramming in this setting.

In comparison, our syngeneic model with C57BL/6J mice bearing YUMM1.7 tumors, Tfap2a knockout alone was sufficient to trigger tumor regression and TLSs formation. The phenotype in this setting was indeed far more dramatic, suggesting the full immunologic consequence of TFAP2A loss requires adaptive immunity.

Together, these two models are complementary rather than redundant. The NSG experiments establish that TFAP2A cell-autonomously suppresses innate immune infiltration in the resistance context; the syngeneic experiments reveal that removing this suppression, in the presence of an intact immune system, is sufficient to trigger organized adaptive antitumor immunity. The logical synthesis is that in patients receiving BRAF/MEK inhibitors, who retain an intact immune system, TFAP2A upregulation simultaneously promotes drug tolerance and actively suppresses the immune response that might otherwise control residual disease.

### BRAF/MEK inhibition undermines antitumor immunity through TFAP2A induction

This interpretation is supported by clinical observations showing that melanoma patients responsive to BRAF/MEK inhibitors initially display increased lymphocytic infiltration, but that this infiltration is not sustained despite continued drug exposure^36–38^. Our syngeneic model recapitulates this dynamic: recurrent tumors become immune-cold over time. We propose that sustained TFAP2A upregulation by BRAF/MEK inhibitors provides a mechanistic explanation for this failure — by progressively creating an immunosuppressive microenvironment and preventing TLSs formation, TFAP2A effectively neutralizes the transient immune activation that follows initial tumor regression. This raises the possibility that BRAF/MEK inhibitors carry an unrecognized immunological cost that limits the durability of response and undermines combination immunotherapy strategies.

### TFAP2A loss triggers TLSs formation, providing a tractable model for studying organized antitumor immunity

A particularly significant finding is that TFAP2A knockout alone is sufficient to initiate TLSs formation. This is notable because the upstream triggers of TLSs in tumors remain incompletely understood, and experimentally inducible TLSs models are rare. The TLSs induced by Tfap2a depletion here recapitulates the canonical architecture including T cell zones with mature dendritic cells adjacent to B cell follicles; therefore, this is a tractable system for dissecting the cellular and molecular events that govern TLSs assembly. This model can be exploited to study how neoantigens are presented by dendritic cells, how antigen-specific T cells are primed and maintained, and how B cells are programmed to produce antitumor antibodies^39–41^. A limitation worth noting is that the TFAP2A-KO TLS model reflects complete, constitutive gene loss rather than the partial or dynamic suppression that might be achieved therapeutically; whether pharmacological attenuation of TFAP2A activity to a lesser degree is sufficient to trigger TLSs in established tumors remains to be tested.

### Macrophage reprogramming as a mechanism linking TFAP2A to the immune microenvironment

Beyond TLSs, TFAP2A loss alone was sufficient to shift macrophage polarization from an M2-like immunosuppressive state toward an M1-like antitumor phenotype. This is mechanistically significant because macrophage polarization is a key determinant of whether the tumor microenvironment supports or suppresses T cell activity^42^. The precise mechanism by which TFAP2A instructs macrophage phenotype is not yet established but is likely indirect. One possibility is that this process is mediated through changes in tumor cell-secreted cytokines, remodeled extracellular matrix, or altered chemokine gradients that modify cell-cell communication between melanoma cells and infiltrating myeloid cells^43,44^. Defining these intermediary signals is an important direction for future work, as they may represent more tractable therapeutic targets than TFAP2A itself.

### Therapeutic implications and targeting strategies

The dual role of TFAP2A, promoting drug resistance while suppressing antitumor immunity, makes it an attractive therapeutic target, as its inhibition would be expected to simultaneously re-sensitize tumors to BRAF/MEK inhibitors and restore immune surveillance. A limitation of directly targeting TFAP2A is that, as a transcription factor, it lacks an obvious enzymatic active site and is therefore difficult to drug with conventional small molecules. We propose two complementary alternative strategies. First, systematic characterization of TFAP2A’s upstream regulators and downstream transcriptional targets may identify nodal proteins that are more amenable to pharmacological inhibition. Second, nanoparticle-mediated delivery of shRNA or CRISPR/Cas9 constructs targeting TFAP2A directly to melanoma cells represents an emerging approach that bypasses the need for a small-molecule binding pocket. Both strategies warrant further preclinical evaluation, particularly in models that combine BRAF/MEK inhibition with immune checkpoint blockade, where the clinical stakes of overcoming immunosuppression are highest.

## Conclusion

In summary, we demonstrate that TFAP2A is induced by BRAF/MEK inhibition and functions as a convergent driver of drug resistance and immune evasion in melanoma. Its roles in the immunocompromised and immunocompetent settings are distinct but mechanistically coherent: TFAP2A cell-autonomously suppresses innate immune infiltration and, in the presence of an intact adaptive immune system, prevents the formation of organized antitumor immune structures. Targeting TFAP2A or its pathway therefore represents a one-stone-two-bird strategy, overcoming targeted therapy resistance while simultaneously unleashing antitumor immunity, and merits prioritization as a therapeutic approach in drug-resistant, immune-cold melanoma.

## Methods

### Cell culture

293T cells or YUMM1.7 cells were cultured in DMEM (Corning, 10-017-CM) medium supplemented with 10% FBS (GeminiBio, 100-106). Patient-Derived-Xenograft (PDX)-derived melanoma cell lines were established in Dr. Heryln’s lab at the Wistar Institute following previous protocols^45^ and the instructions of the kit (Miltenyi Biotec #130-095-929). Dissociated cells were cultured in MCD153 medium (Sigma-Aldrich, F8105-500ML) supplemented with 2% FBS (GeminiBio, 100-106), 20% Leibovitz’s L-15 medium (Corning, 10-045-CV) and 1.65mM CaCl2 for establishment of melanoma cell lines.

### CRISPR/Cas9 cloning for genome editing

All CRISPR single guide RNAs (sgRNAs) were designed and cloned following previous protocols^46^. Briefly, designed single strand oligos were annealed and ligated into a lentiCRISPRv2-blasticidin backbone (Addgene, 98293) cut with BsmBI-v2 (NEB, R0739S). Ligation products were transformed into One Shot Stbl3 competent cells (Invitrogen, C737303), plated onto LB-agar plates containing 50 mg/mL ampicillin and cultured overnight at 37°C incubator. The next day, individual clones were picked up and inoculated as mini culture. Using the kit (Qiagen, 12123), plasmid DNAs were isolated from individually picked clones and submitted to the Wistar Genomic Facility for Sanger sequencing to validate the success of the sgRNA cloning.

### Lentivirus production and selection of pooled or single knockout cells

293T cells were seeded onto 10 cm dishes on Day 1. LentiCRISPRv2 plasmid carrying sgRNA/Cas9, pCMV-VSV-G plasmid (Addgene, 8454) and pCMV-dR8.2 dvpr (Addgene, 8455) plasmid were mixed altogether and transfected to 293T cells with LT1 transfection reagent (Mirus Bio LLC, MIR2300) on Day 2. Fresh media were changed on Day 3. Supernatant was filtered using 0.45 mm syringe filters (Corning, 975421) on Day 4 to infect melanoma cells. Polybrene (Millipore, TR-1003-G) was added to enhance the efficiency of virus infection. On Day 5, Blasticidin (Life Technology, A1113903) was added to select the infected cells carrying CRISPR/Cas9 cassettes. To get single knockout cells, we seeded 1000 single cells onto 10 cm dish, cultured them for 3-4 weeks until cell colonies formed. We then scraped individual colonies under the microscope carefully, transfer the scrape cells onto 12-well plates, and expanded them.

### Colony formation assay

Cells were counted, seeded onto 6-well plates (10,000 cells per well) and then treated with indicated drugs. After 1-2 weeks, cell colonies were stained with crystal violet solution (0.05% cystal violet (w/v), 1% formaldehyde (v/v), 1% Methanol (v/v) in 1x PBS) for 30 min at room temperature. Stained wells of plates were then imaged, and area percentage of crystal violet stained colonies were quantified by the Wistar Image Facility.

### Western blotting

Cells were lysed in RIPA buffer (CST, 9806) containing protease/phosphatase inhibitor cocktail (CST, 5872) and centrifuged at 12,000 rpm to remove cell debris. Supernatant were transferred to new Eppendorf tubes. Protein concentrations were quantified using Pierce BCA protein assay kit (Thermo Scientific, 23228). Samples were boiled for 15 min in 4X NuPAGE LDS sample buffer (Invitrogen, NP008) containing 100 mM DTT to denature proteins. 20 μg proteins were loaded into wells of 4-12% NuPAGE Bis-Tris mini protein gels (Invitrogen; NP0321BOX). Proteins were separated based on their size in the gel and then transferred onto PVDF membrane (Bio-Rad, 1620177). We blocked the membrane using blocking buffer (Li-Cor, 927-70001) for 1 hour at room temperature. Primary antibodies with 1:2000 dilution against BRAF (CST, 14814), TFAP2A (CST, 3215), or ACTIN (CST, 3700S) were added to the membrane and incubated overnight at 4°C. Anti-rabbit secondary antibody with IRDye 800CW (Li-Cor, 926-32213) or anti-mouse secondary antibody with 680RD (Li-Cor, 926-68070) were added and incubated for 2 hours at room temperature. Final signals were visualized using Li-Cor Odyssey imaging machine.

### Histology and Immunohistochemistry

We euthanized the mice and collected desired tumors or tissues and then fixed them overnight in 4% formalin (v/v) overnight. Fixed tissues were then switched to cold PBS and submitted to the Wistar Histology Facility for paraffin embedding and sectioning. Hematoxylin and eosin (H&E) stained slides and unstained slides were obtained from the histology facility. We then performed immunohistochemistry (IHC) on unstained slides. Briefly, we deparaffinized the unstained slides in 100% xylene for three times. We then hydrated the slides in serial of ethanol (v/v) 100%, 95%, 90% and 70%. After boiling the slides for 10 min in 1x antigen unmasking solution buffer (Vector Laboratories, H-3300-250), we followed ImmPRESS Excel Amplified HRP Polymer Staining kit (Vector Laboratories, MP-7601) for the IHC staining. Primary antibody against F4/80 (CST, 70076S) was diluted 1:200 in PBST buffer and incubated with slides overnight at 4°C on a shaker. We then followed the protocol from the same kit to get the final signal. The slides were then counterstained with hematoxylin and dehydrated in serial ethanol. Slides were then immersed in 100% xylenes three times and sealed with Cytoseal 60 (Thermo Scientific, 8310-16) for long term storage. All images were scanned, uploaded and organized by the Wistar Image Facility with the aid of the Concentriq software.

### Immunofluorescent co-staining

Desired tissues were collected and fixed in 4% formalin (v/v) overnight. Tissues were then submitted to the Wistar Histology Facility for paraffin embedding and sectioning. Slides were deparaffinized in 100% xylene for three times and then hydrated in serial of ethanol (v/v) 100%, 95%, 90% and 70%. Slides were boiled for 10 min in 1x antigen unmasking solution buffer (Vector Laboratories, H-3300- 250) for antigen retrieval. Next, we followed the ImmPRESS Excel Amplified HRP Polymer Staining kit (Vector Laboratories, MP-7601). Antibody against HMB45 (Dako, M063401-2), F4/80 (70076), Cd3 (CST, 40932), Cd20 (CST, 69874) or Cd11c (CST, 64675) were incubated with the slides overnight. The next day, slides were washed with PBST and incubated with secondary antibody for 1 hour if primary antibody is not conjugated with fluorescent signal. Slides were then incubated with DAPI for nuclei staining and finally sealed with prolong gold antifade mountant medium (Thermo Fisher Scientific, P36930). Images were captured using confocal microscopy at the Image facility of Wistar Institute.

### Flow cytometry

The whole tumor was dissociated into single cells using tumor dissociation kit (Miltenyi Biotec, 130-095-929) following the manufacturer’s instructions. Cells were then stained with antibodies against F4/80 (Biolegend, 157310) and Cd11b (Biolegend, 101212). FlowJo was used to analyze the raw data.

### Trichrome staining

Tumors were collected and fixed in 4% formalin (v/v) overnight. Tissues were submitted to the histology facility at the Wistar Institute for paraffin embedding and sectioning. Unstained slides were used for trichrome staining following the protocol of the trichrome stain kit (Abcam, ab150686). Stained slides were scanned for images at the Wistar image facility.

### Bulk RNA sequencing

mRNA was isolated using the RNeasy Plus Mini Kit (Qiagen, C74134). We then used the NEBNext® rRNA Depletion Kit v2 (NEB, E7400) to deplete ribosome RNA (rRNA) and NEBNext® Ultra™ II Directional RNA Library Prep Kit (NEB, 7760) to prepare RNA sequencing libraries. Prepared RNA libraries were sequenced by 75 + 75 paired-end reading using NextSeq2000 (Illumina).

The RNA-seq raw data was aligned to UCSC hg38 reference genome using STAR (v2.7.11)^47^. Gene-level expression quantification was computed using HTSeq (v2.0.4)^48^ for differential expression analysis with DEseq2 (v1.42)^49^. The raw counts were further normalized to TPM (transcripts per kilobase million) values for gene set enrichment analysis (GSEA, v4.0.2)^50^ from Gene ontology (GO) (v2023.2)^51^ dataset.

### Chromatin ImmunoPrecipitation Sequencing (ChIP-seq)

ChIP-seq was performed as desbribed previously^52^. Briefly, we culture the cells and cross-link proteins with DNAs with 1% formaldehye. Cell lysates were sheared with sonication and pulled down with specific antibodies as the following: IgG control (CST, 2729S), TFAP2A (Proteintech, 13019-3-AP). Pulled down DNA fragments were then washed and eluted. Prepared DNA libraries using a kit (NEB, 7760) were sequenced by 75 + 75 paired-end reading using NextSeq550 (Illumina). ChIP-seq data were processed starting with quality trimming of raw fastq files using Trim_Galore (v0.6.10) (https://github.com/FelixKrueger/TrimGalore) to remove low-quality and adapter-contaminated reads. High-quality reads were then aligned to the UCSC hg38 reference genome using BWA-MEM (v.0.7.18)^53^. Duplicate reads were marked using GATK’s MarkDuplicates tool^54^. Peak calling was conducted with MACS2^55^, and differential binding analysis was performed using DiffBind (v3.14.0) (https://bioconductor.org/packages/release/bioc/vignettes/DiffBind/inst/doc/DiffBind.pdf) with a fold change cutoff greater than 1 and FDR below 0.05. Motif analysis was carried out using HOMER(v5.1)^56^ to identify motifs within these high-confidence binding regions and comparing them with known motifs in the JASPAR database to validate the findings. Visualization of differential binding regions was achieved using deepTools2(3.5.6)^57^.

### Single nuclei RNA sequencing

Tumors were collected and used for isolation of single nuclei (Illumina, 20132795). 20,000 single nuclei per sample were subjected to cDNA preparation using the 10x single cell RNA sequencing kit (10X genomics,1000285). Reads alignment was performed using Cell Ranger v8.0.0 (10x Genomics). All scRNA-seq expression matrices were processed using the Seurat v5^58^. Cells were filtered out if they had fewer than 500 detected genes, more than 50,000 unique molecular identifiers (UMIs), or over 20% of the reads mapped to mitochondrial genes. Doublets were identified and removed using DoubletFinder v2.0.4^59^. After quality control, the samples were integrated and normalized using SCTransform^60^. Principal component analysis (PCA) was performed, and the top 10 components were retained based on the elbow point of the standard deviation plot for downstream analysis. Cell types were annotated manually based on canonical markers.

### Tumor growth

Human melanoma cells were trypsinized, washed with PBS, and resuspended cold Matrigel diluted with cold PBS (v/v, 50%). One million cells were subcutaneously injected into the lower right flank of each mouse. Tumor was measured twice every week using a digital caliper. At indicated time points, tumors were collected, and flash frozen in liquid nitrogen or processed into Formalin-Fix-Paraffin-Embedded (FFPE) blocks to be used for further analysis.

### Drug Treatments

For *in vitro* drug treatment, melanoma cells were seeded onto 10cm dishes, and treated with Dabrafenib/Trametinib for 7 days. Cell lysates were then collected and subjected to western blot to check the protein level of TFAP2A.

For *in vivo* trial, Dabrafenib (150 mg/kg) and Trametinib (1.5 mg/kg) were administered orally to mice via chow (Bio-Serv, S7581).

### Statistics

We work with Dr. Qin Liu at the Wistar Institute bioinformatic core for statistics. p values are shown in figures or supplementary tables among groups.

## Supporting information

Table for p value

## Data availability

The bulk RNA-seq, ChIP-seq, and single-cell RNA-seq raw sequencing data generated in this study have been deposited in the Gene Expression Omnibus (GEO) under separate accession numbers and will be made publicly available upon publication. Reviewer access is available upon request.

## Acknowledgements

We thank the Wistar Core Facilities (Animal, Histology, Imaging and Genomic sequencing). This work is supported by NIH grants P50 CA261608, P01 CA114046, U54 CA224070, R01 CA238237, R01 CA240362, R01 CA258113, R01 CA259295, P30 CA010815, R01-CA227115 and the Dr. Miriam and Sheldon G. Adelson Medical Research Foundation.

## Author contributions

H.M. and M.H. conceived and designed the experiments. H.M., V.Y., N.S., M.X., M.D., and K.D. performed the experiments. H.M., V.Y., N.S., M.X., M.D., K.D., and J.L.S. analyzed and interpreted the data. Y.C. did the bioinformatic analysis. H.M. and J.L.S. wrote the manuscript under the supervision of M.H.

## Conflict of interests

The authors declare no conflict of interests.

## Supplementary Figures and legends

## Supplementary Figures and legends

**Supplementary Figure 1.**
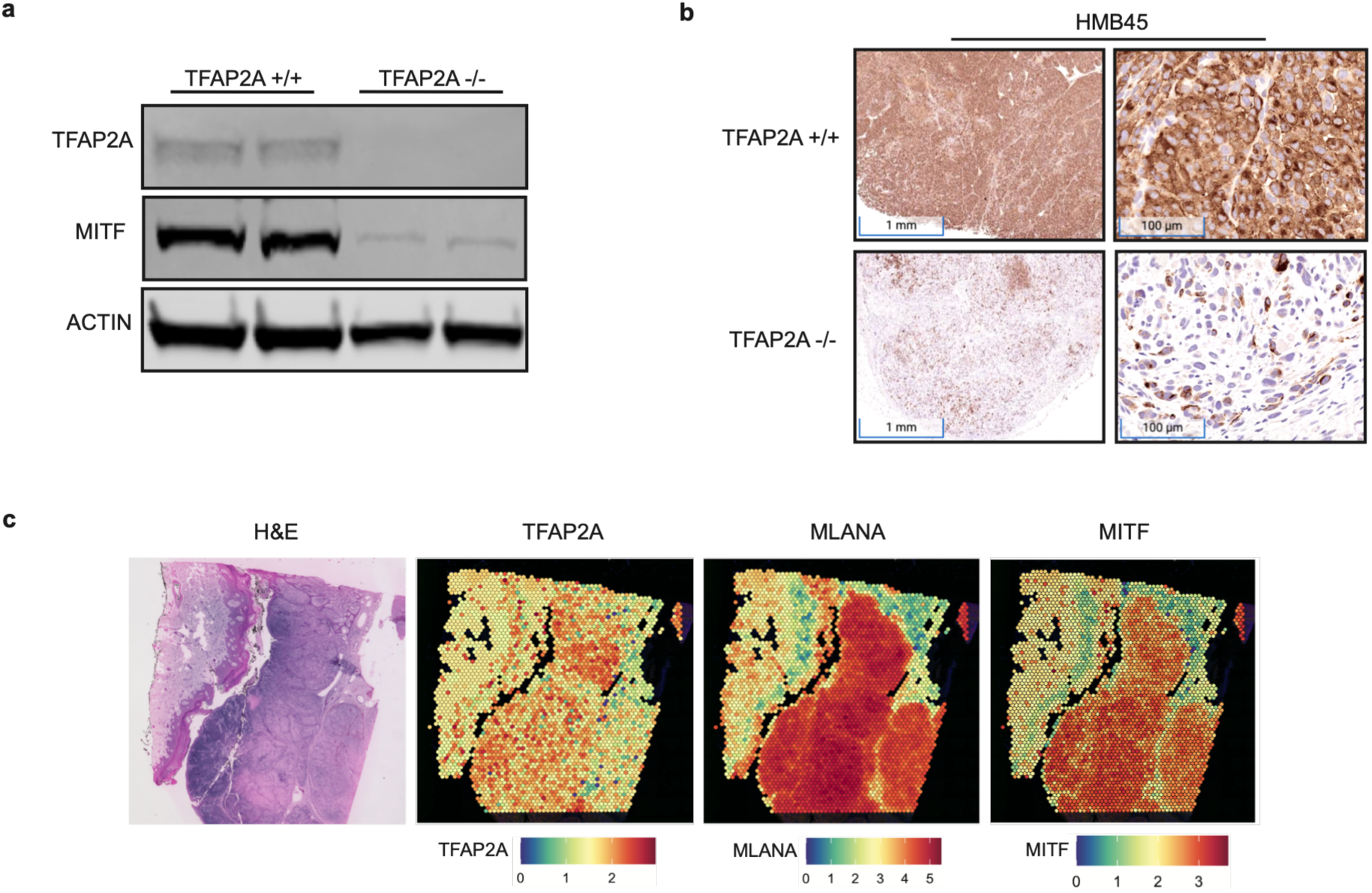
TFAP2A regulates pigmentation genes. (**a**) Western blot shows that MITF expression is downregulated in TFAP2A-KO cells. (**b**) TFAP2A-KO melanomas show less pigmentation marker, HMB45. (**c**) Correlation analysis from public spatial single cell RNA sequencing dataset reveals that TFAP2A spatially correlates well with the expression of MLANA and MITF, which are the pigmentation molecules.

**Supplementary Figure 2.**
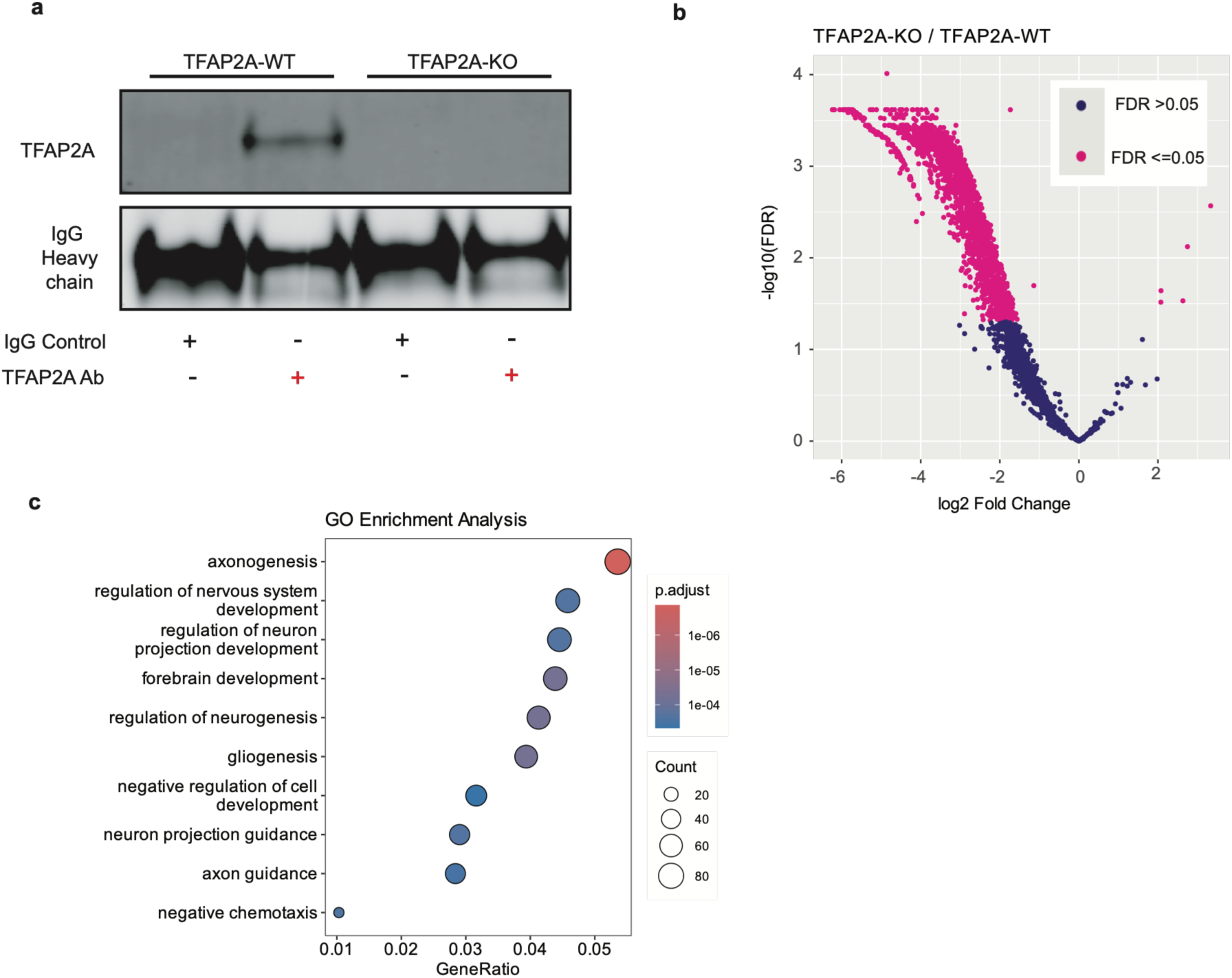
ChIP-seq to identify direct transcriptional targets of TFAP2A. (**a**) Co-Immunoprecipitation (CoIP) shows antibody against TFAP2A successfully pull down the target protein TFAP2A. (**b**) ChIP-seq using the antibody in (a) successfully pulled down gene fragments which are annotated and presented as volcano plot. (**c**) GSEA analysis using the annotated genes from (b) show TFAP2A targets are enriched in neurogenesis pathways.

**Supplementary Figure 3.**
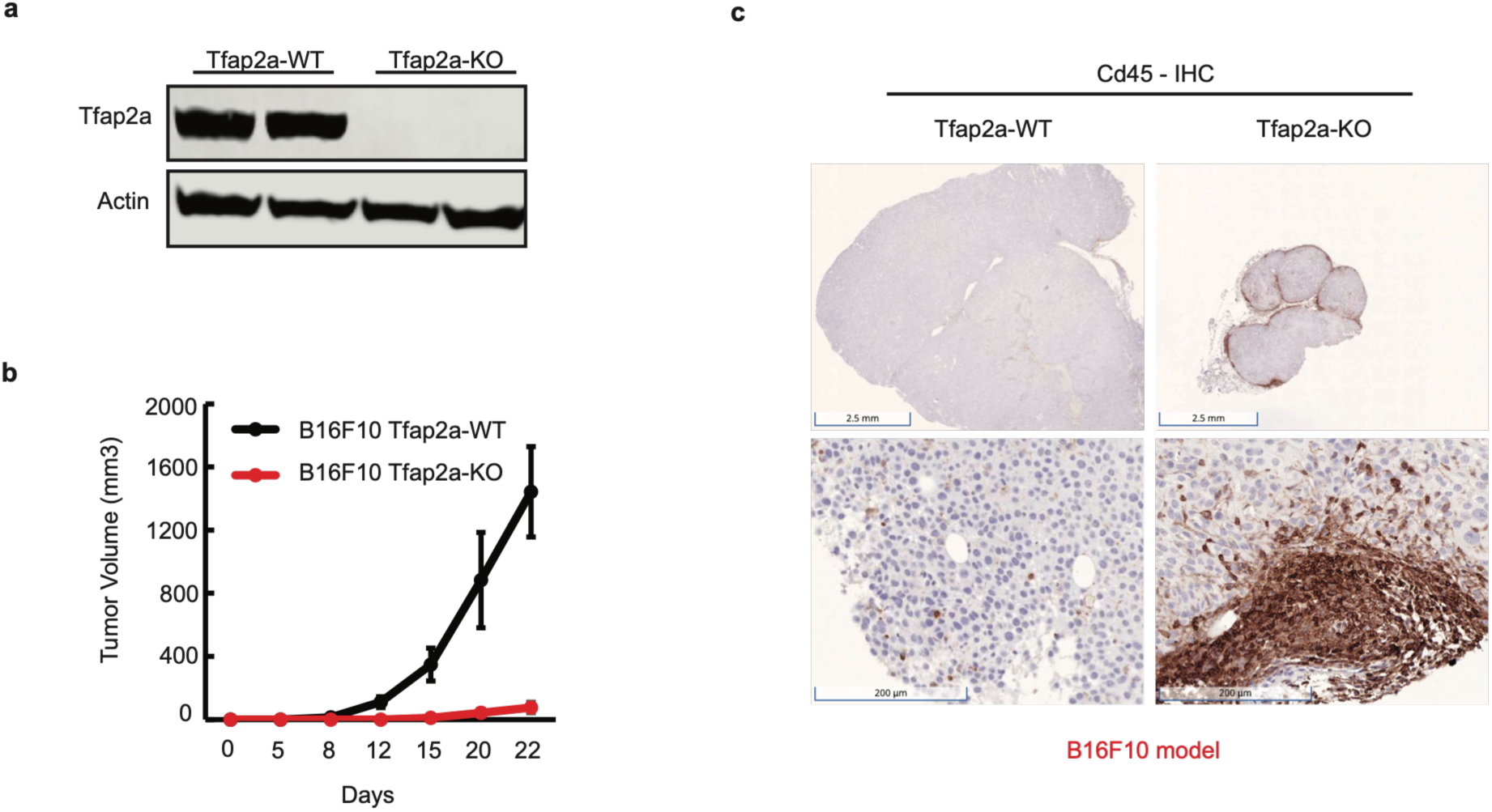
Tfap2a knockout impairs tumor growth and promotes formation of immune clusters in B16F10 melanoma model. (**a**) Western blot confirms the protein loss of Tfap2a in B16F10 knockout cells. (**b**) Tumor growth curve of B16F10-Tfap2a-WT or B16F10- Tfap2a-KO melanomas in C57BL/6J mice (n=5). (**c**) Immunohistochemistry (IHC) staining of Cd3+ T cells in Tfap2a-WT or Tfap2a-KO melanomas. Brown color indicates the positive staining of Cd3+ cells.

**Supplementary Figure 4.**
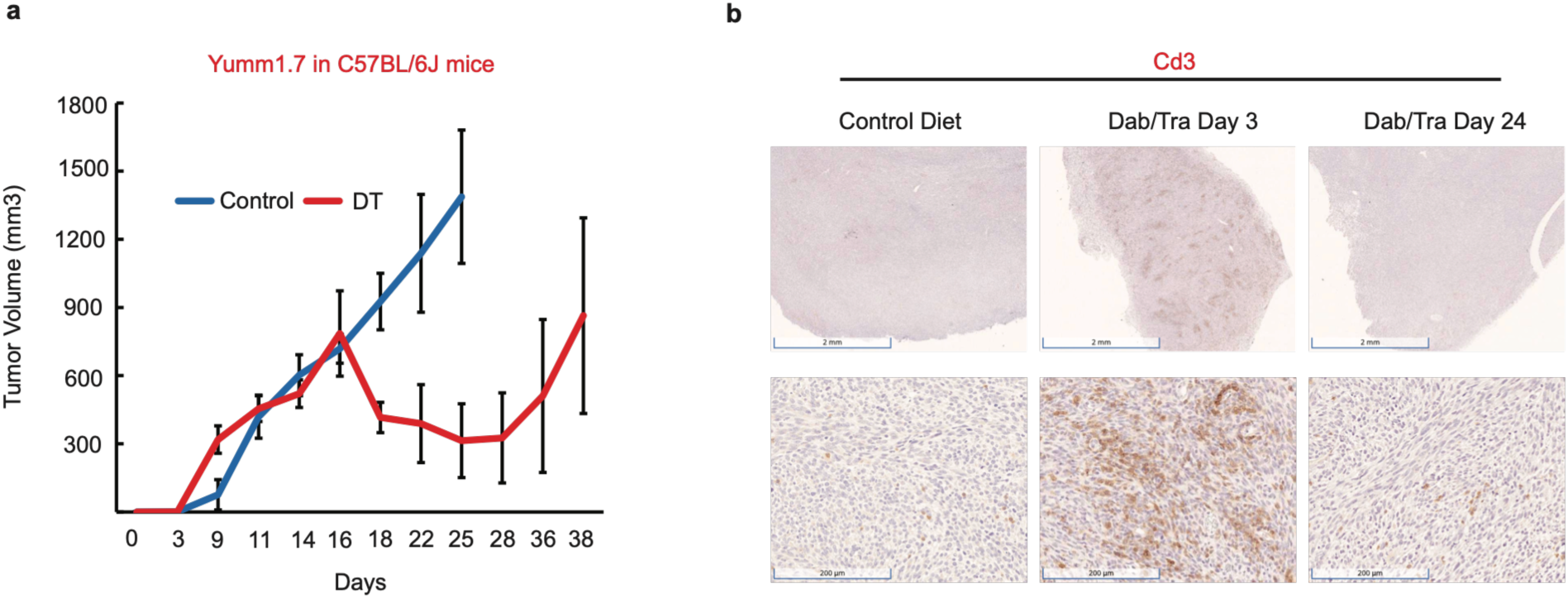
Recurrent melanomas show immune-cold feature. (**a**) Yumm1.7 cells were subcutaneously injected into NSG mice for tumor growth curve. Dabrafenib/Trametinib (D/T) chow or control chow were given on day 16. Red curve shows the Yumm1.7 melanomas respond to BRAF/MEK inhibitors (D/T) initially, shrink robustly and quickly develop relapse. (**b**) Immunohistochemistry (IHC) staining of Cd3+ T cells in Yumm1.7 melanomas treated with Dabrafenib/Trametinib (Dab/Tra) for different period of times (Day 0, Day 3 and Dady 24). Brown color indicates the positive staining of Cd3+ cells.

**Supplementary Figure 5.**
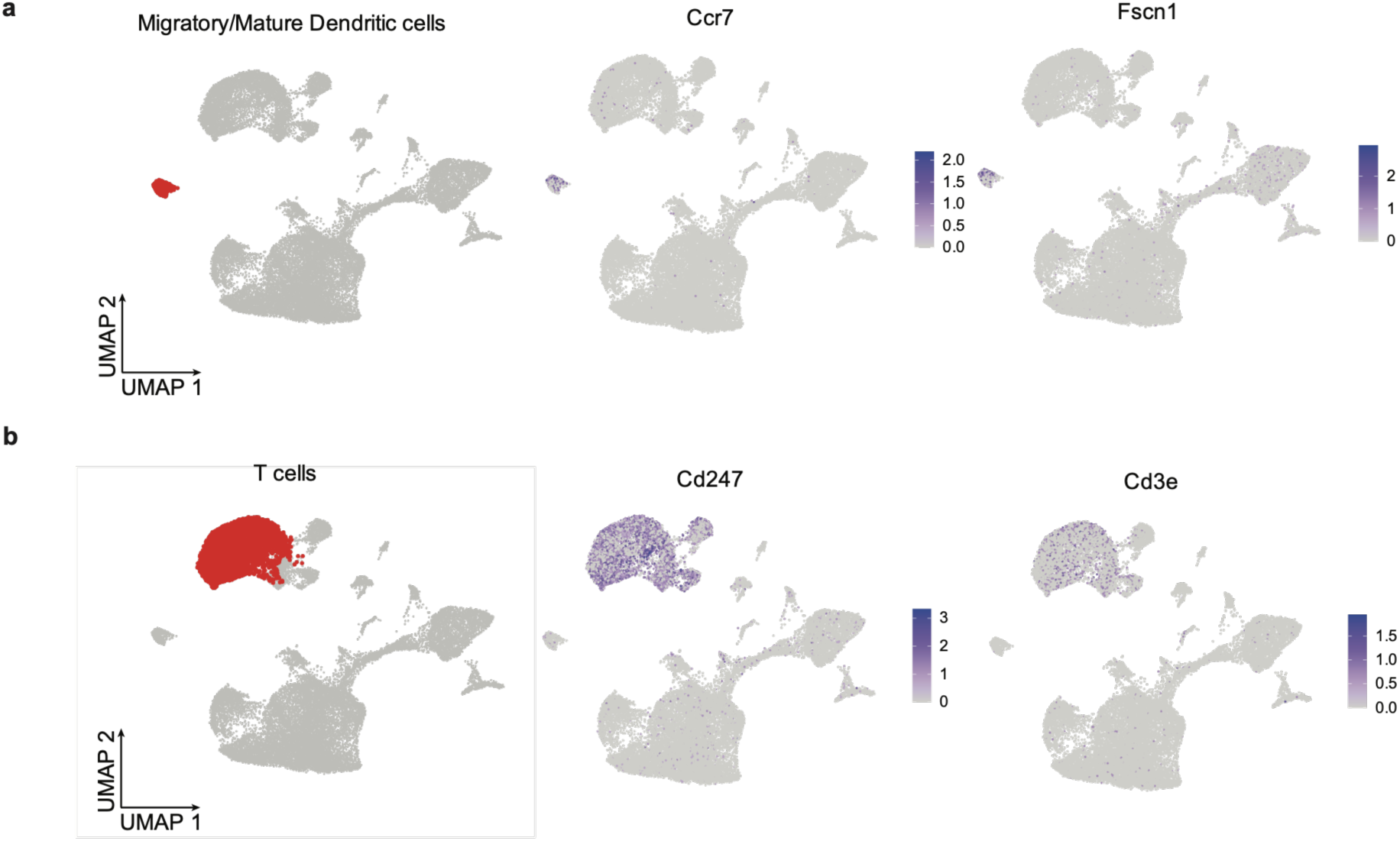
Sing cell RNA-seq annotation of Dendritic cells and T cells. (**a**) Dendritic cell subset within Yumm1.7 melanomas. Ccr7 and Fscn1 are markers for mature dendritic cells. (**b**) T cell subset. Cd247 and Cd3e are markers for T cells.

## Notes

### Competing Interest Statement

The authors have declared no competing interest.

