## Supplementary material for "TFAP2A links drug resistance to antitumor immunity": Table for p value

**Figure 3e (p values)**

**Methods:** ANOVA with Tukey's multiple comparisons test were used to compare the tumor volume
at day 44 among 4 groups.

| Tukey’s multiple comparison test | significance | Adjusted p value |
|---|---|---|
| TFAP2A-WT vs TFAP2A-KO | Yes. | 0.001 |
| TFAP2A-WT+Dab/Tra vs TFAP2A-KO+Dab/Tra | Yes. | 0.0417 |
| TFAP2A-WT vs TFAP2A-WT+Dab/Tra | No. | 0.0610 |
| TFAP2A-KO vs TFAP2A-KO+Dab/Tra | No. | 0.8086 |
